# Enhanced wort fermentation with *de novo* lager hybrids adapted to high ethanol environments

**DOI:** 10.1101/204198

**Authors:** Kristoffer Krogerus, Sami Holmström, Brian Gibson

## Abstract

Interspecific hybridization is a valuable tool for developing and improving brewing yeast in a number of industry-relevant aspects. However, the genomes of newly formed hybrids can be unstable. Here, we exploited this trait by adapting four brewing yeast strains, three of which were *de novo* interspecific lager hybrids with different ploidy levels, to high ethanol concentrations in an attempt to generate variant strains with improved fermentation performance in high-gravity wort. Through a batch fermentation-based adaptation process and selection based on a two-step screening process, we obtained eight variant strains which we compared to the wild-type strains in 2L-scale wort fermentations replicating industrial conditions. The results revealed that the adapted variants outperformed the strains from which they were derived, and the majority also possessed several desirable brewing-relevant traits, such as increased ester formation and ethanol tolerance, as well as decreased diacetyl formation. The variants obtained from the polyploid hybrids appeared to show greater improvements in fermentation performance. Interestingly, it was not only the hybrid strains, but also the *S. cerevisiae* parent strain, that appeared to adapt and showed considerable changes in genome size. Genome sequencing and ploidy analysis revealed that changes had occurred both at chromosome and single nucleotide level in all variants. Our study demonstrates the possibility of improving *de novo* lager yeast hybrids through adaptive evolution by generating stable and superior variants that possess traits relevant to industrial lager beer fermentation.

**Importance:** Recent studies have shown that hybridization is a valuable tool for creating new and diverse strains of lager yeast. Adaptive evolution is another strain development tool that can be applied in order to improve upon desirable traits. Here we apply adaptive evolution to newly created lager yeast hybrids by subjecting them to environments containing high ethanol levels. We isolate and characterize a number of adapted variants, which possess improved fermentation properties and ethanol tolerance. Genome analysis revealed substantial changes in the variants compared to the original strains. These improved variants strains were produced without any genetic modification, and are suitable for industrial lager beer fermentations.

## Introduction

Yeast breeding and hybridization has in recent years been shown to be a promising tool for developing and improving brewing yeast in a number of industry-relevant aspects (1–4). These include improving fermentation rates, sugar use, and aroma compound production. However, the genomes of newly formed hybrids tend to be unstable, and these may undergo substantial structural changes after the hybridization event (5–9). As yeast is commonly reused for multiple consecutive fermentations in industrial breweries (even over 10 times depending on brewery), it is vital that the genomes of any newly developed yeast strains remain stable to ensure product quality during industrial use. Yeast encounter a range of challenges during brewery fermentations, such as low oxygen availability, osmotic stress, CO_2_ accumulation, nutrient limitation and ethanol toxicity (10), which may contribute to faster changes in the genome size (11, 12). Yeast strains with larger genomes (i.e. with a higher ploidy level) in particular have been shown to show greater changes in genome size during such conditions (9, 11–15). This may especially cause concerns when a rare mating approach is used for hybridization (3, 16). However, genome stabilisation can be achieved, for example, by growing newly formed hybrids for 30–70 generations under fermentative conditions (2, 9). Phenotypic changes may occur though during the genome stabilisation process, altering the properties of the original hybrid (6).

To influence the changes occurring during the stabilisation process, an adaptive or experimental evolution approach could be applied. Studies have demonstrated that adaptive evolution can be used, for example, to obtain strains with increased tolerance to ethanol (14, 17–19), high-gravity wort (20, 21), lignocellulose hydrolysates (22), and extreme temperatures (23, 24), as well as improved consumption of various sugars (25–27). Numerous recent studies utilizing experimental evolution have also provided valuable information on what genetic changes take place in the yeast strains during adaptation to various stresses (5, 8, 14, 15, 22–24, 28, 29). Evolution experiments with yeast hybrids have shown that various changes may occur during adaptation, including partial loss of one of the parental sub-genomes, loss of heterozygosity and selection of superior alleles, and the formation of fusion genes following translocations (5, 8, 22, 28). Studies have also revealed that the ploidy of the yeast has an effect on adaptability, with tetraploid strains appearing to adapt more rapidly than diploid strains (12, 15, 30). Taking this into consideration, we sought to not only stabilize, but simultaneously adapt a range of our newly created lager yeast hybrids to conditions normally encountered during brewery fermentations. One of the main stresses brewing yeast encounter during the fermentation process is that of increasing ethanol concentrations (10). Particularly, as interest from the industry towards very high gravity fermentations (i.e. those with wort containing over 250 g extract / L, resulting in beer with alcohol contents above 10% (v/v)) has increased in recent years (31).

In this study, we therefore exposed 3 *de novo* lager yeast hybrids of different ploidy and a common *S. cerevisiae* ale parent strain to 30 consecutive batch fermentations in media containing 10% ethanol in an attempt to retrieve variant strains with increased tolerance to ethanol. Following the adaptation stage, isolates were screened and selected based both on their ability to ferment wort sugars efficiently in the presence of ethanol, and their ability to ferment high gravity wort. Eight variant strains, two from each of the original strains, were ultimately selected and compared in 2L-scale wort fermentations. Analysis of the fermentations and resulting beers revealed that all variants appeared to outperform the original strains during fermentation. Furthermore, the majority of the variants produced beers with higher concentrations of desirable aroma-active esters and lower concentrations of many undesirable aroma compounds, such as higher alcohols and diacetyl. The genomes of the variant strains were also sequenced, and genome analysis revealed that changes had occurred both at chromosome and single nucleotide level.

## Results

Three different *de novo* lager yeast hybrids, generated in previous studies by our lab (3, 32), along with a *S. cerevisiae* ale parent strain (common to all three hybrids) were subjected to the adaptation process (Table 1). The ploidy of the interspecific hybrids varied from around 2.4N to 4N. The four yeast strains, referred to as Y1-Y4 according to Table 1, were grown for 30 consecutive batch fermentations in two different media containing 10% ethanol in an attempt to generate ethanol-tolerant variants with improved fermentation properties (Figure 1A). The first medium, M1, contained 2% maltose as a fermentable carbon source, while the second, M6, contained 1% maltose and 1% maltotriose. These sugars were chosen as they are the main sugars in all-malt wort. Over the 30 fermentations, approximately 130 to 161 yeast generations were achieved depending on yeast strain and growth media (Figure 2). The optical density at the end of each batch fermentation increased from around 2.5- to over 10-fold depending on the yeast strain, suggesting adaptation to the high ethanol concentration (Figure 2). Isolates from each adaptation line were obtained after 10, 20 and 30 fermentations, by randomly selecting the fastest growing colonies on agar plates containing solidified versions of the same adaptation media (Figure 1B).

**Table 1.** Yeast strains used in the study.

| Working code | Species | Information | Measured ploidy | Source |
| --- | --- | --- | --- | --- |
| Y1 | *S. cerevisiae* | A *S. cerevisiae* ale strain (VTT-A81062) | 1.95 ($\pm 0.15$) | VTT Culture Collection |
| Y1_M1 | *S. cerevisiae* | Variant obtained from Y1. Isolated after 30 fermentations from media M1, replicate A. | 2.02 ($\pm 0.21$) | Isolated in this study |
| Y1_M6 | *S. cerevisiae* | Variant obtained from Y1. Isolated after 30 fermentations from media M6, replicate A. | 3.64 ($\pm 0.17$) | Isolated in this study |
| Y2 | *S. cerevisiae* × *S. eubayanus* | A tetraploid interspecific hybrid between strain Y1 and the *S. eubayanus* type strain VTT-C12902. Known as 'Hybrid H1' in the source study. | 4.03 ($\pm 0.25$) | (32) |
| Y2_M1 | *S. cerevisiae* × *S. eubayanus* | Variant obtained from Y2. Isolated after 30 fermentations from media M1, replicate B. | 3.47 ($\pm 0.26$) | Isolated in this study |
| Y2_M6 | *S. cerevisiae* × *S. eubayanus* | Variant obtained from Y2. Isolated after 30 fermentations from media M6, replicate B. | 3.57 ($\pm 0.31$) | Isolated in this study |
| Y3 | *S. cerevisiae* × *S. eubayanus* | A triploid interspecific hybrid between strain Y1 and the *S. eubayanus* type strain VTT-C12902. Known as 'Hybrid B3' in the source study. | 2.98 ($\pm 0.22$) | (3) |
| Y3_M1 | *S. cerevisiae* × *S. eubayanus* | Variant obtained from Y3. Isolated after 30 fermentations from media M1, replicate B. | 3.03 ($\pm 0.27$) | Isolated in this study |
| Y3_M6 | *S. cerevisiae* × *S. eubayanus* | Variant obtained from Y3. Isolated after 30 fermentations from media M6, replicate B. | 3.19 ($\pm 0.23$) | Isolated in this study |
| Y4 | *S. cerevisiae* × *S. eubayanus* | An interspecific hybrid containing DNA from strain Y1, *S. cerevisiae* WLP099 (White Labs Inc.) and the *S. eubayanus* type strain VTT-C12902. Known as 'Hybrid T2' in the source study. | 2.38 ($\pm 0.24$) | (32) |
| Y4_M1 | *S. cerevisiae* × *S. eubayanus* | Variant obtained from Y4. Isolated after 30 fermentations from media M1, replicate A. | 2.27 ($\pm 0.25$) | Isolated in this study |
| Y4_M6 | *S. cerevisiae* × *S. eubayanus* | Variant obtained from Y4. Isolated after 20 fermentations from media M6, replicate B. | 2.27 ($\pm 0.24$) | Isolated in this study |

**Figure 1.**
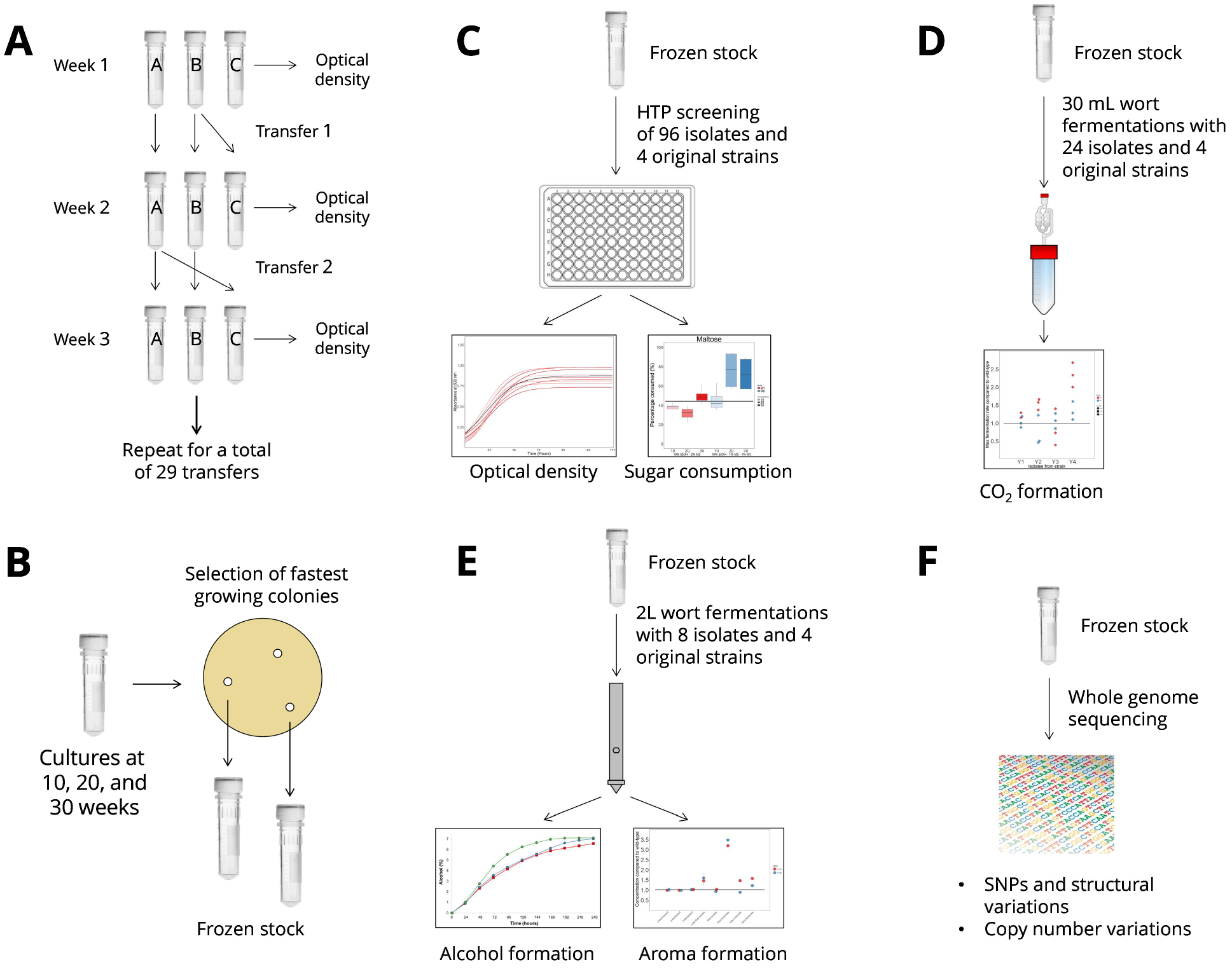
Experimental overview. (**A**) 30 consecutive batch fermentations were carried out with four different yeast strains and two different ethanol-containing media in duplicate adaptation lines. (**B**) An initial set of isolates were obtained by selecting fast-growing colonies on solidified versions of the adaptation media. (**C**) High-throughput screening of all the isolates was performed in a malt extract-based screening media containing ethanol. The best-performing isolates were chosen for further screening based on the maltose and maltotriose consumption. (**D**) Small-scale wort fermentations were performed with selected isolates to ensure they were able to ferment wort efficiently and perform in media without exogenous ethanol. (**E**) 2L-scale wort fermentations replicating industrial conditions were performed with 8 variant strains (Table 1) and vital aroma compounds of the resulting beers were analysed. (**F**) The genomes of the 8 variant strains were sequenced and compared to those of the wild-type strains. For more information, see the Materials & Methods section.

**Figure 2.**
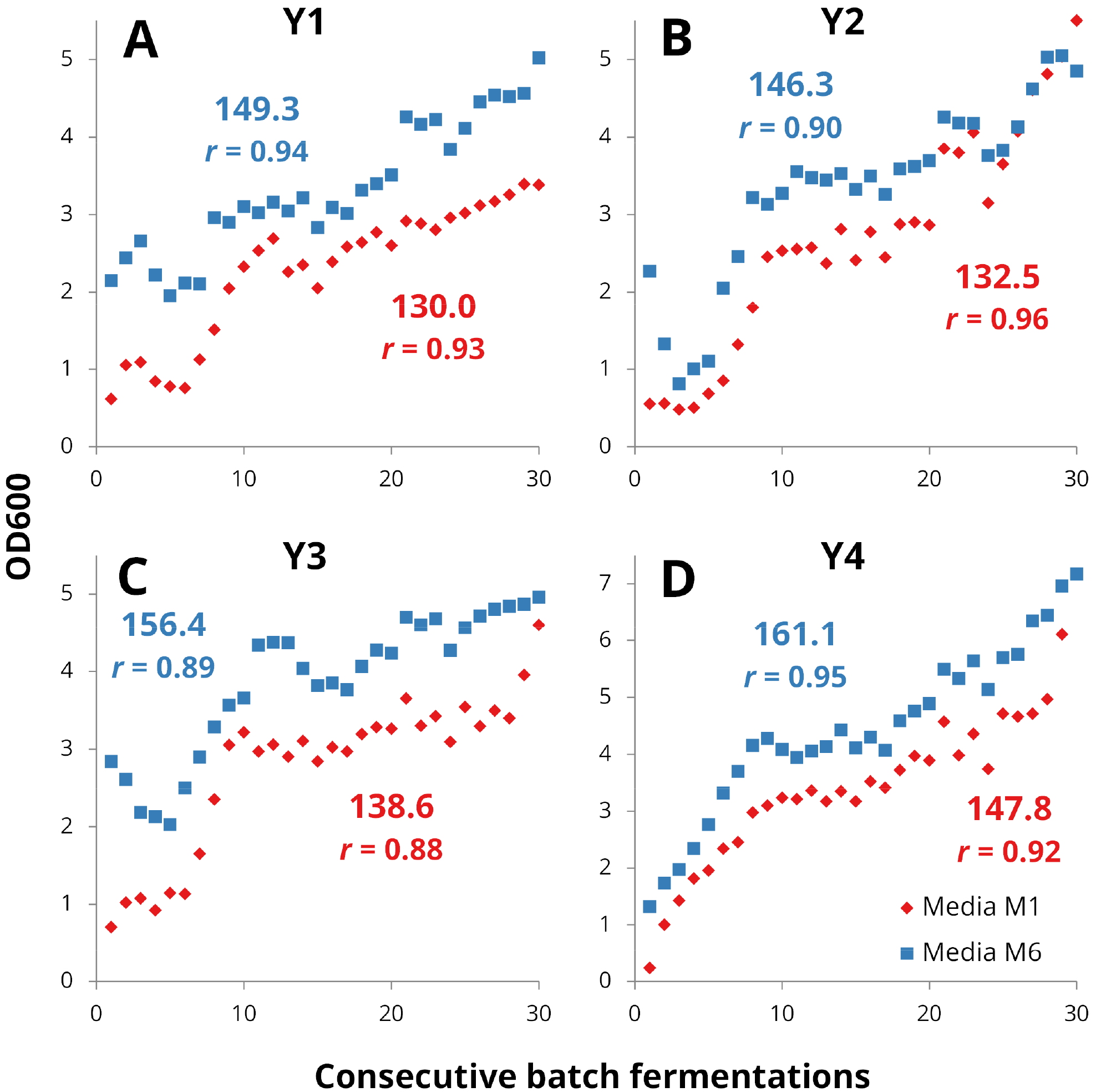
The optical densities at the end of each consecutive batch fermentation with yeast strains (**A**) Y1, (**B**) Y2, (**C**) Y3, and (**D**) Y4 in the two different ethanol-containing media (*red diamonds*: Media M1 (10% ethanol, 2% maltose); *blue squares*: Media M6 (10% ethanol, 1% maltose, 1% maltotriose)). The cumulative number of yeast generations after the 30^th^ batch fermentation and Pearson’s correlation coefficient (*r*) between the optical densities and number of consecutive fermentations is presented above in blue and below in red for the yeast grown in Media M6 and M1, respectively.

### Screening of isolates reveals improved sugar consumption and fermentation rates

The 96 isolates that were obtained from the adaptation fermentations were then subjected to high-throughput screening in a malt-based media containing ethanol and sorbitol (Figure 1C). The ethanol and sorbitol were added to replicate the stresses the yeast is exposed to during brewery fermentations. The majority of the variant strains grew similarly to the wild-type strains, and all strains were able to reach stationary growth phase during the 144 hour cultivation period (Figure S1 in Supplementary material). As the objective was to select variant strains with enhanced Termentation rate, rather than enhanced growth, we also monitored the sugar concentrations in the media at three time points.

There were considerable differences in the amounts of maltose and maltotriose consumed between the wild-type and variant strains after 144 hours of fermentation (Figure 3). There was no obvious pattern between the consumption of the different sugars, the isolation time points (i.e. the amount of consecutive batch fermentations), and the two different adaptation media among the variants strains. In many cases, the largest consumption of both maltose and maltotriose was observed with variants that had been isolated after 30 batch fermentations. However, with variants obtained from yeast strain Y2, the average maltose and maltotriose consumption of variants obtained after 30 batch fermentations was lower than those isolated at earlier stages (Figure 3B). Nevertheless, the variant strain derived from Y2 with the highest maltose consumption was obtained after 30 batch fermentations. Several variant strains from all four wild-type strains (Y1-Y4) showed higher sugar consumption than the wild-type strains. In total, 83% of the variants consumed more maltose, and 60% consumed more maltotriose than the wild-type strains during the screening fermentations. Interestingly, all variants that consumed more maltotriose than the wild-type strains also consumed more maltose. Excluding maltotriose from the adaptation media did not appear to have any negative effect on maltotriose consumption in the variants, as the variant strains derived from Y2, Y3, and Y4 with the highest maltotriose consumption were obtained from the adaptation media lacking maltotriose. 6 variants per wild-type strain were selected for further screening in small-scale wort fermentations, based on the highest sugar consumptions and the requirement that they were derived from separate adaptation lines and isolation time points.

**Figure 3.**
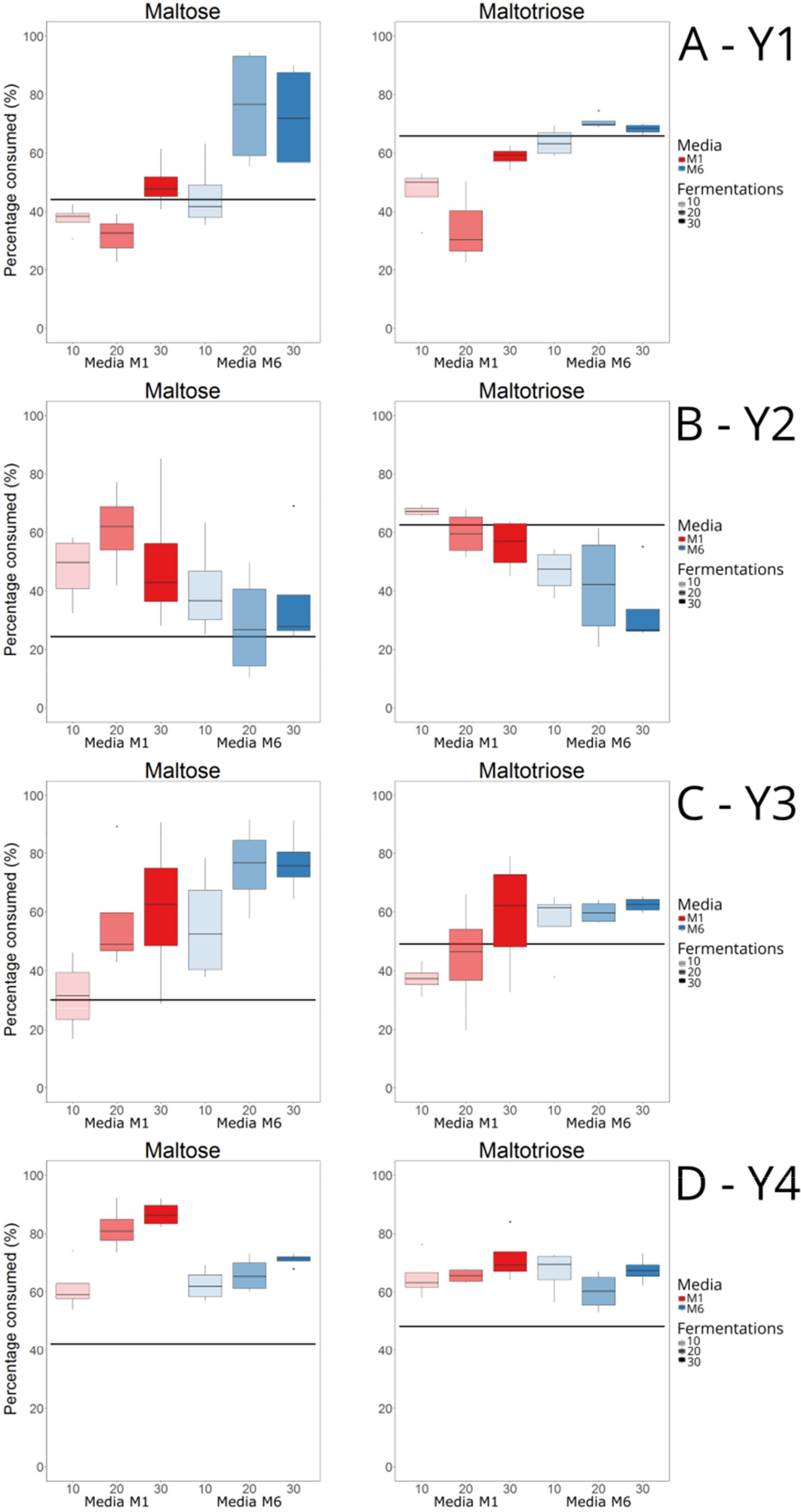
The percentage of maltose and maltotriose consumed by isolates (selected after 10, 20 and 30 consecutive batch fermentations) of the yeast strains (**A**) Y1, (**B**) Y2, (**C**) Y3, and (**D**) Y4 after 144 hours of fermentation in the screening media (6.2% malt extract, 10% sorbitol, 5% ethanol) during high-throughput screening in 96-well plates. The black line depicts the amount of sugar consumed by the wild-type strain (average calculated from 12-16 replicate fermentations). For each of the three isolation points (10, 20 and 30 consecutive batch fermentations), four isolates were selected per yeast strain per media (a total of 24 isolates per parent strain). Three replicate fermentations were carried out for each isolate.

Small-scale wort fermentations were used as a final screening step to ensure that the selected variants were also able to ferment wort efficiently and perform in media without exogenous ethanol (Figure 1D). A 15 °P high gravity wort, i.e. a wort similar to what is used in the brewing industry, was used for the fermentations. They revealed that 17 out of the 24 tested variants outperformed the wild-type strains from which they were derived (Figure 4) in regards to the maximum fermentation rate that was observed. Of these 17 variants, 13 also reached a significantly higher (*p* < 0.05 as determined by two-tailed Student’s t-test) final alcohol level after the 9 days of fermentation (data not shown). Variants which had been adapted in the media containing 2% maltose as the sugar source (Media M1) appeared to outperform those obtained from the media containing both 1% maltose and maltotriose (Media M6). One isolate per yeast strain and media (for a total of 8 isolates) were selected for more thorough characterization in 2L-scale wort fermentations. These isolates (listed in Table 1) were selected based on the highest fermentation rates, and those that had undergone 30 batch fermentations were also preferentially selected.

**Figure 4.**
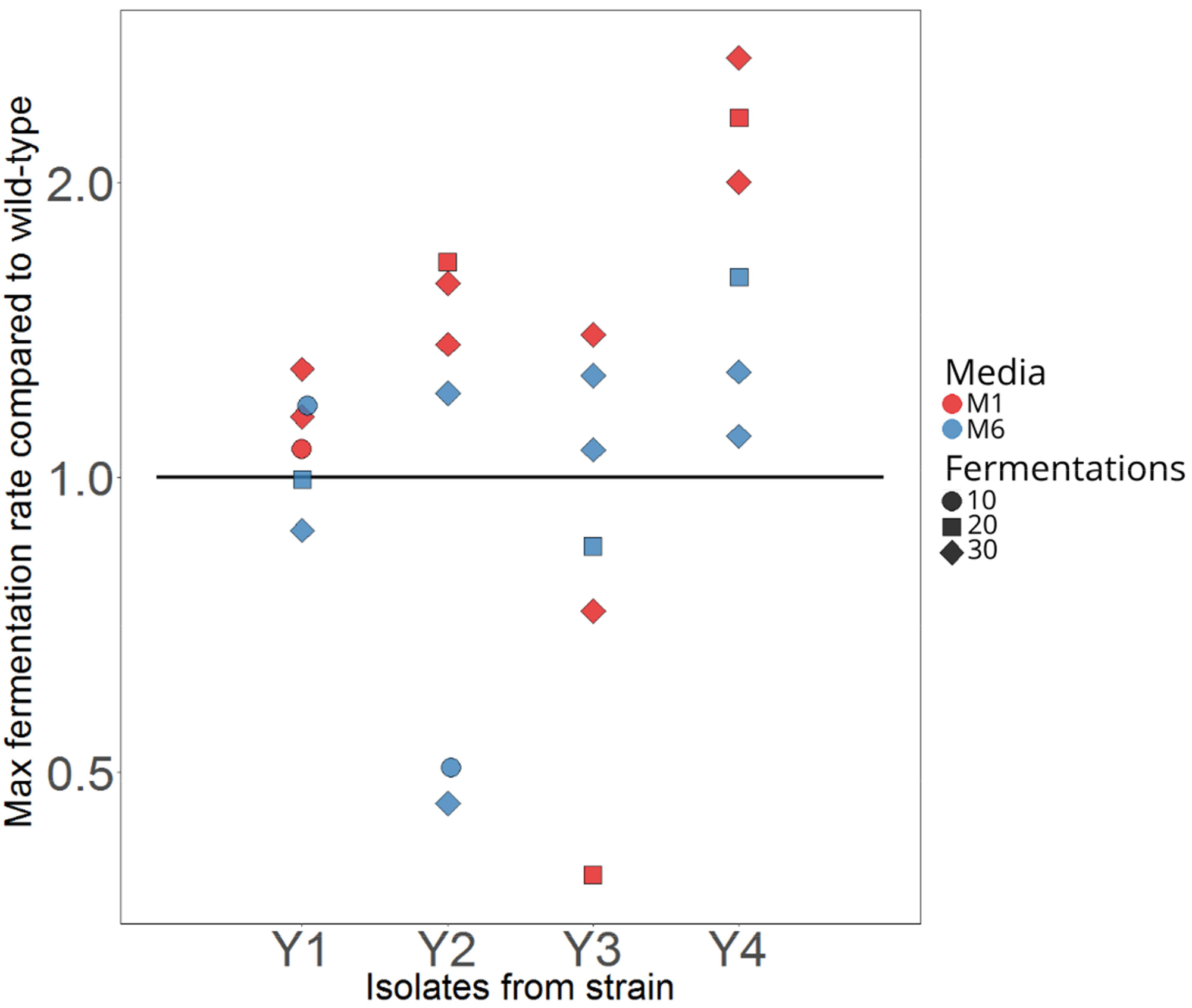
The maximum fermentation rate of 24 isolates compared to their wild-type strains during small-scale fermentations in 15 °P all-malt wort at 15 °C. Isolates were selected based on sugar consumption during high-throughput screening and were from the two different growth media (M1 and M6 in *red* and *blue*, respectively), and three different isolation points (10, 20 and 30 consecutive batch fermentations, with *circles*, *squares* and *diamonds*, respectively). Duplicate fermentations were carried out for each isolate.

### Enhanced performance confirmed in 2L-scale wort fermentations

In order to examine how the variant strains (Table 1) perform in a brewery environment, 2L-scale tall-tube fermentations were carried out in high-gravity 15 °P all-malt wort at 15 °C (Figure 1E). These conditions were chosen to replicate those of industrial lager fermentations. All eight variant strains appeared to outperform their respective wild-type strains during these fermentations (Figure 5). Time-points after which a significant difference (*p* < 0.05 as determined by Student’s t-test) was observed between the variant and the wild-type strain are marked with arrows in the plot. The largest differences in fermentation compared to the wild-type strains were observed with the variants of Y2, i.e. the tetraploid interspecific *S. cerevisiae* × *S. eubayanus* hybrid, and those of Y3, i.e. the triploid interspecific *S. cerevisiae* × *S. eubayanus* hybrid. For most strains, differences between variant and wild-type strains seemed to appear after approximately 48 hours of fermentation. Before this time point, it is mainly the monosaccharides that are consumed from the wort and the alcohol level is still below 2% (v/v) (Figure S2 in Supplementary material). The sugar profiles during fermentation also revealed that improved maltose consumption appears to be one of the main causes for the increased fermentation rate of the variant strains. Three of the variants, derived from strains Y2 and Y4, only showed a difference compared to the wild-type strain late in fermentation. These observations suggest that the observed differences may be due to the variant strains possessing an enhanced ability to ferment maltose and maltotriose or to tolerate increasing ethanol concentrations in the wort.

**Figure 5.**
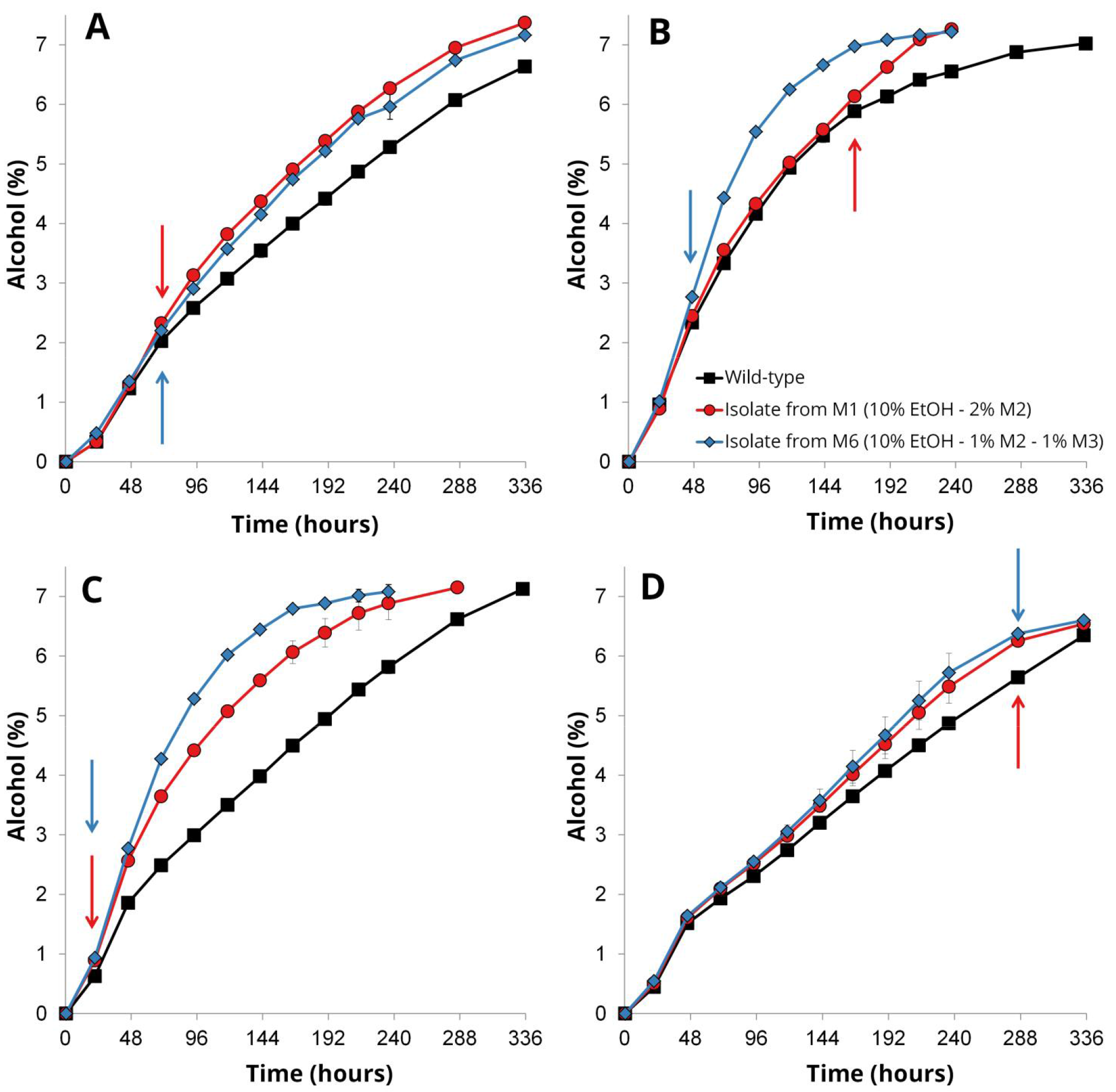
The alcohol content (% ABV) in the beers fermented at 2L-scale from 15 °P wort at 15 °C with wild-type (*black squares*) and variant (*red circles* and *blue diamonds*) strains derived from yeast strains (**A**) Y1, (**B**) Y2, (**C**) Y3, and (**D**) Y4. Values are means from two independent fermentations and error bars where visible represent the standard deviation. Arrows indicate the time-point after which a significant difference was observed between the variant and wild-type strain as determined by two-tailed Student’s t-test (*p* < 0.05).

We also wanted to compare the aroma profiles of the beers produced with the variant strains with those produced with the wild-type strains, to ensure that the adaptation process hadn’t introduced any negative side effects to the resulting beer. Genetic hitchhiking is common during adaptive evolution (33), and here we only screened and selected for an increased fermentation rate. Analysis of the aroma-active higher alcohols and esters in the beers revealed that the variant strains, in general, produced equal or lower amounts of unwanted higher alcohols, while equal or higher amounts of desirable esters compared to the wild-type strains (Figure 6). The concentrations of 3-methylbutyl acetate, possessing a banana-like flavour (34), and ethyl esters, possessing fruity and apple-like flavours (34), in particular appeared to increase in the variant strains. We also monitored the concentrations of diacetyl, an important unwanted off-flavour in lager beer fermentations (35), and results revealed that five out of eight variant strains had produced significantly lower concentrations of diacetyl than the wild-type strains, while the other three produced concentrations that were equal to the wild-type strains. Hence, results revealed that the adaptation process had not only yielded variant strains with improved fermentation performance in wort, but also, inadvertently, strains that produced more desirable aroma profiles. In addition, all eight variant strains appeared genetically stable over 80 generations (Figure S3 in Supplementary material).

**Figure 6.**
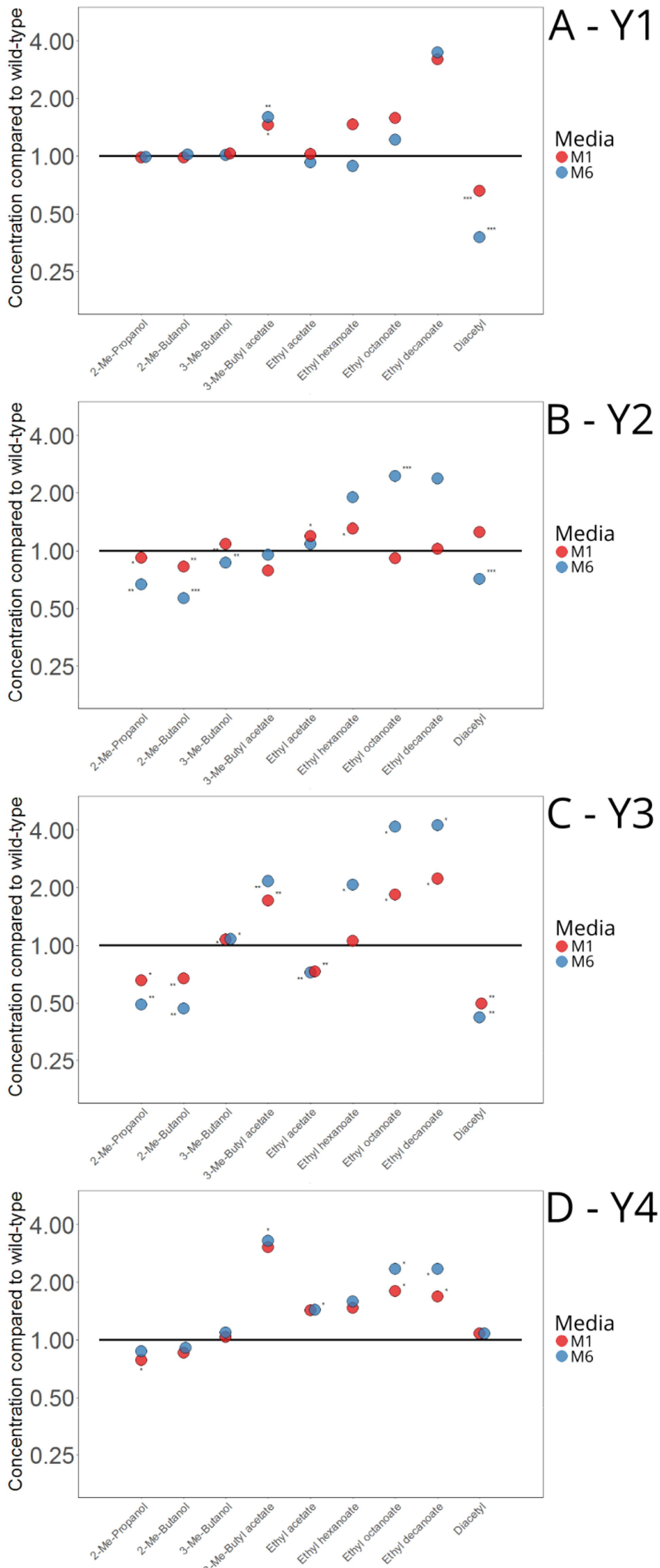
The concentrations of nine yeast-derived aroma compounds in the beers fermented with the variant strains relative to those fermented with the wild-type strains (**A**) Y1, (**B**) Y2, (**C**) Y3, and (**D**) Y4. Values are means from two independent fermentations and asterisks depict a significant difference in the variant compared to the wild-type as determined by two-tailed Student’s t-test (* *p* < 0.05; ** *p* < 0.01; *** *p* < 0.001). Me: methyl.

### Ethanol tolerance and accumulation capacity of variant strains

As the variant strains (Table 1) were derived from repeated exposure to high ethanol concentrations and they performed better particularly towards the end of high-gravity wort fermentations, we wanted to test and compare their ethanol tolerance and accumulation capacity to that of the wild-type strains. All strains were able to grow on YPM agar supplemented with 9% ethanol (v/v), but differences in growth were revealed on YPM agar supplemented with 11% ethanol (v/v) (Figure 7). The variant strains derived from Y4 in particular, showed improved growth at 11% ethanol compared to the wild-type strain (Figure 7D). For the variants derived from the other strains (Y1-Y3), there were no or less obvious differences in the ability to grow in the presence of 11% ethanol. For strain Y2, the variant (Y2_M1) derived from adaptation media M1 (10% ethanol and 2% maltose) also appeared to grow better than the wild-type strain at 11% ethanol (v/v) (Figure 7B). The ethanol accumulation capacity, which measures both the osmo- and ethanol tolerance of a strain, was significantly higher for both variant strains of Y2 and Y4 compared to their wild-type strains, while no significant differences were observed for strain Y3. Surprisingly, the ethanol accumulation capacities of the variants of Y1 were significantly lower than the wild-type strain, despite both variant strains appearing to grow slightly better on YPM agar supplemented with 11% ethanol (Figure 7A).

**Figure 7.**
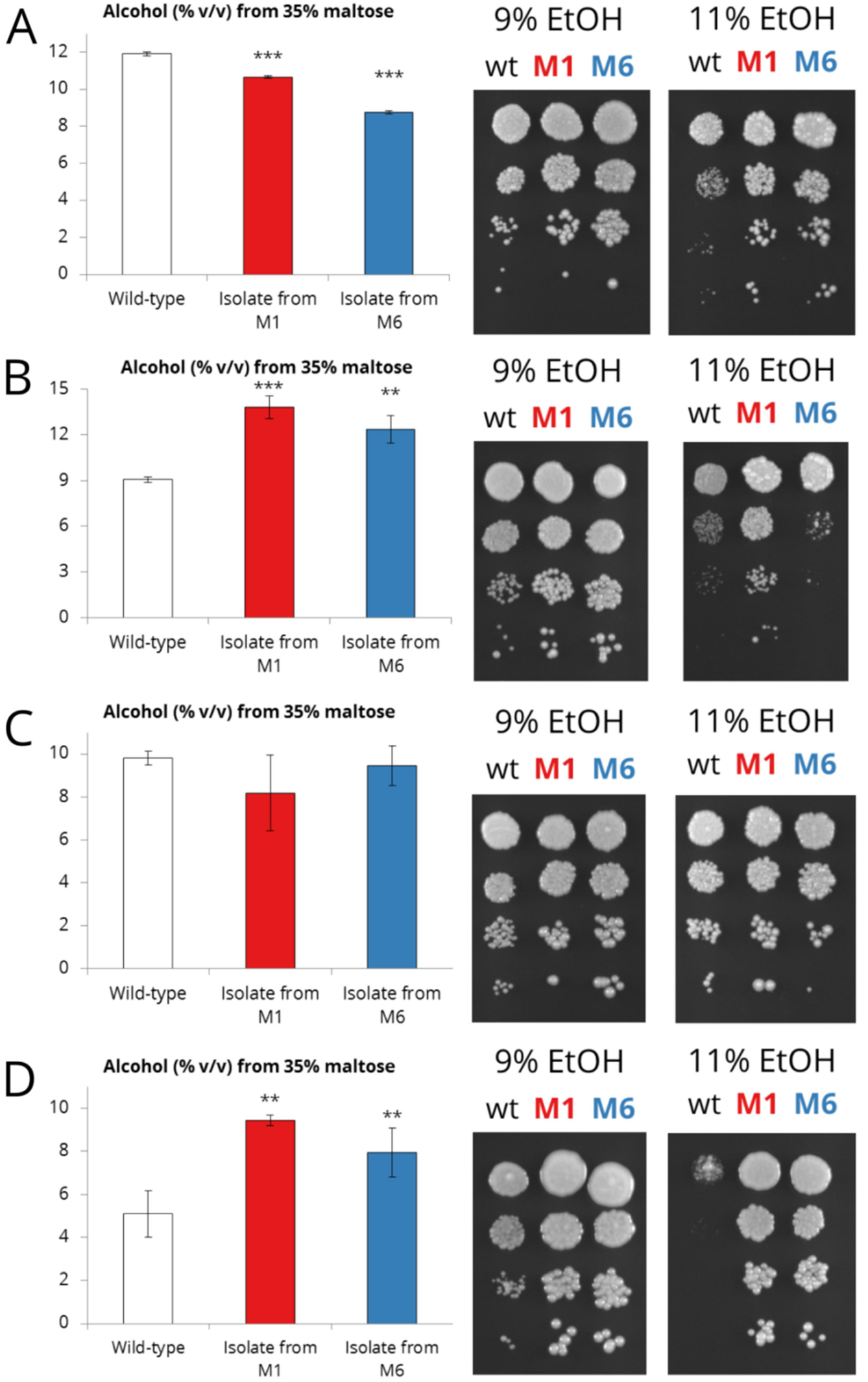
The ethanol accumulation capacity from 35% maltose and the ability to grow on media containing 9% and 11% ethanol of the wild-type (wt) and variant strains derived from yeast strains (**A**) Y1, (**B**) Y2, (**C**) Y3, and (**D**) Y4. Values are means from two independent cultures, error bars where visible represent the standard deviation, and asterisks depict a significant difference in the variant compared to the wild-type as determined by two-tailed Student’s t-test (* *p* < 0.05; ** *p* < 0.01; *** *p* < 0.001).

### Sequencing reveals large-scale changes in genomes

In order to investigate what genetic changes had occurred in the variant strains during the adaptation process, whole genome sequencing and estimation of ploidy by flow cytometry was performed (Figure 1F). Ploidy analysis revealed that relatively large changes in genome size had occurred for many of the variant strains (Table 1). The genome of the variant (Y1_M6) derived from the diploid *S. cerevisiae* strain Y1 and adaptation media M6 had almost doubled in size, while the genomes of both variants (Y2_M1 and Y2_M6) derived from the tetraploid interspecies hybrid Y2 had decreased by approximately 0.5N. Smaller changes were observed in the genome sizes of the variants derived from the triploid and diploid interspecies hybrids Y3 and Y4.

Whole genome sequencing of the wild-type and variant strains (average coverage ranged from 152× to 1212×) revealed both chromosome gains and losses across all variant strains (Figure 8). As indicated by the ploidy analysis, the largest changes in chromosome copy numbers were observed in the variant derived from strain Y1 and adaptation media M6 (Y1_M6), where the majority of the chromosomes were now present in two extra copies. The variants derived from interspecies hybrids (Y2-Y4) had, on average, gained 1.7 and lost 3.2 chromosomes. A greater amount of chromosome copy number changes were also observed in the variants derived from the polyploid hybrids (6.5, 5.5, and 2.5 for Y2, Y3, and Y4, respectively). In regards to the two sub-genomes of the hybrid variants, there were significantly more (*p* < 0.05) chromosome gains in the *S. cerevisiae* sub-genome (average of 1.5 per variant) compared to the *S. eubayanus* sub-genome (average of 0.17 per variant). In the *S. cerevisiae* sub-genome there was no significant difference between the amount of chromosome gains and losses (average of 1.17 per variant). In contrast, the *S. eubayanus* sub-genome had experienced significantly more (*p* < 0.003) chromosome losses (average of 2 per variant) than gains. Common chromosome copy number changes were seen in several variants, as the *S. cerevisiae*-derived chromosomes VII and XIV were amplified in four and six variants, respectively, while the *S. eubayanus*-derived chromosome VII had been lost in four variants (Figure 8).

**Figure 8.**
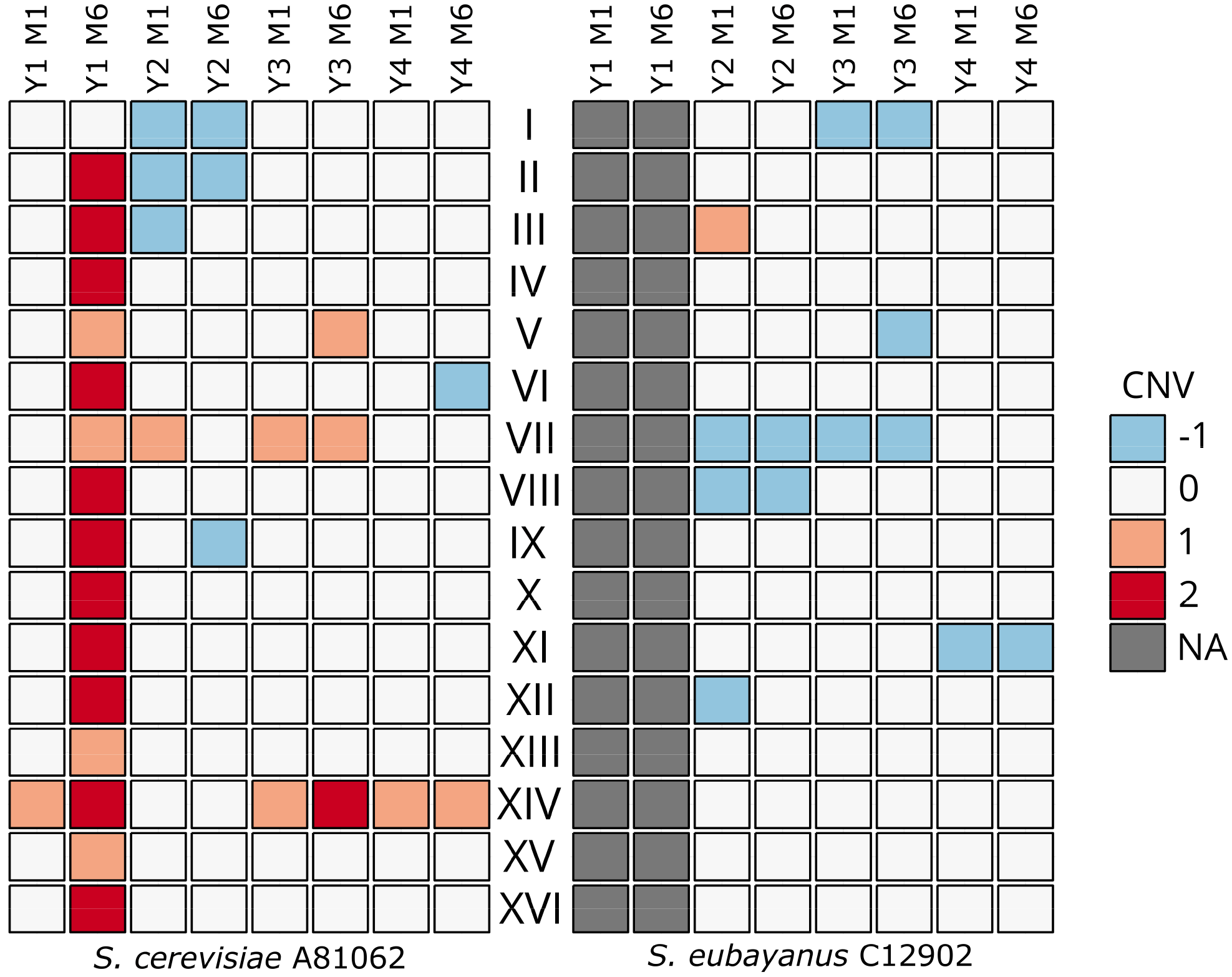
Chromosome copy number variations (CNV) in the *S. cerevisiae* A81062 (left) and *S. eubayanus* C12902 (right) sub-genomes of the variant strains compared to the wild-type strains.

The genomes of the variant strains varied not only at chromosome level, as several unique single nucleotide polymorphisms (SNP), insertions and deletions (Indel) were also observed. A total of 109 unique mutations were identified in the eight variant strains (Table S1 in Supplementary material). Of these 64.2% were intergenic, 8.3% were synonymous, and 27.5% were non-synonymous. In addition, 21% of the mutations were hemi- or homozygous. The non-synonymous mutations caused both amino acid substitutions and frameshift mutations (Table 2 and Table S1 in Supplementary material), and at least one was present in all variant strains. Interestingly, non-synonymous mutations in three genes (*IRA2*, *HSP150*, and *MNN4*) were found in multiple variants. In the case of *IRA2*, an inhibitory regulator of the RAS-cAMP pathway (36) which contained non-synonymous mutations in three of the variants, both the *S. cerevisiae* and *S. eubayanus* orthologues were affected. In addition to the unique mutations that were observed in the variant strains, several of the variant strains had undergone loss of heterozygosity in large regions of several *S. cerevisiae*-derived chromosomes (Figures S4-15 in Supplementary material). The left arms of chromosomes X and XII, as well as the right arm of XV, for example, were affected in multiple variants. No unique translocations or complex structural variations were identified in the variant strains.

**Table 2.**
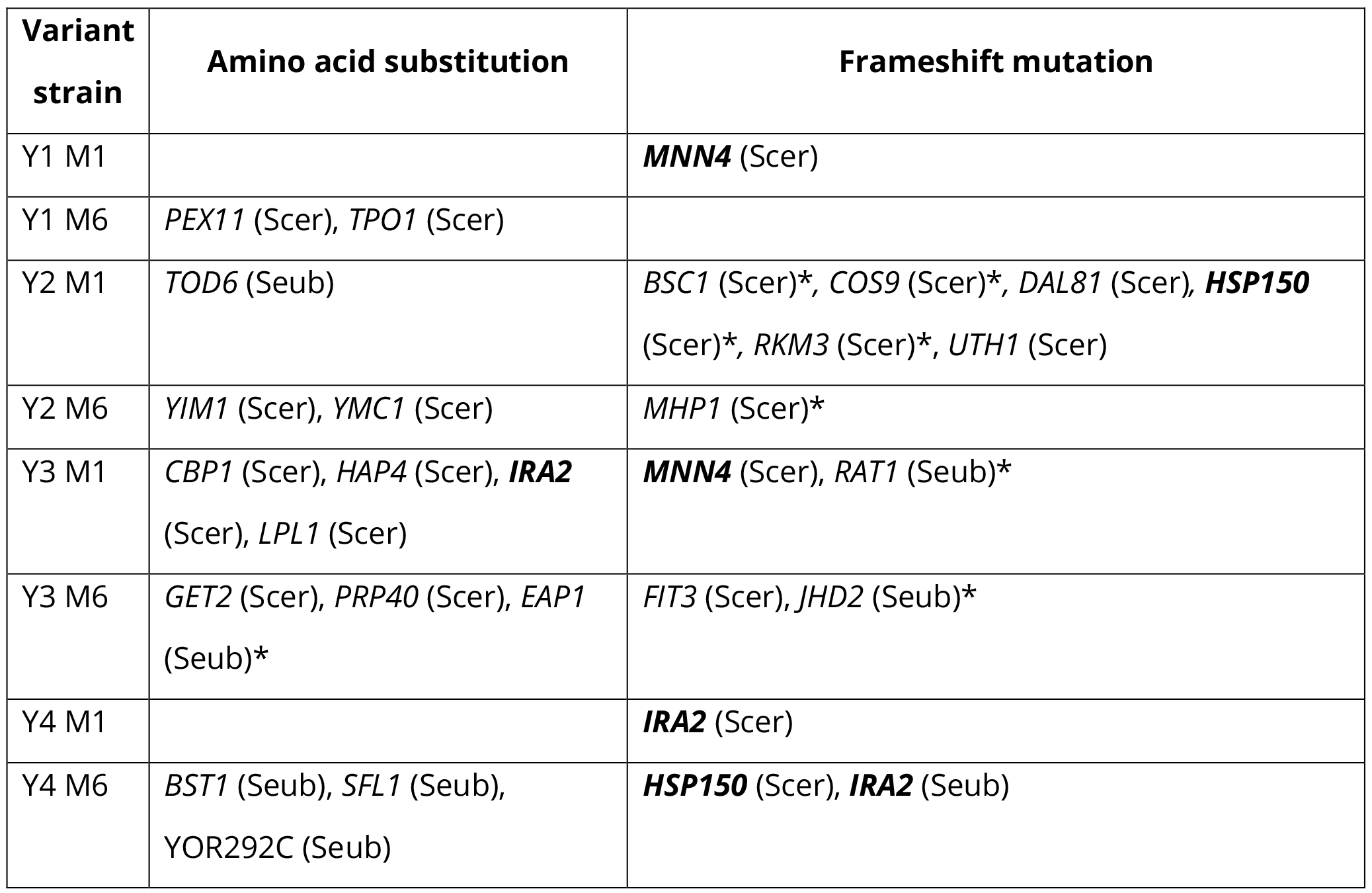
Non-synonymous mutations discovered in the variant strains. Genes that were affected by mutations in several different variants are **in bold**. Whether the *S. cerevisiae* (Scer) or *S. eubayanus* (Seub) orthologue was affected is indicated in parenthesis after the gene name. An asterisk (*) denotes whether the mutation was either homo- or hemizygous. Positions and nucleotide changes of mutations are listed in Table S1 in Supplementary material.

## Discussion

The beer market and industry is driven by an increasing demand for more diverse beer flavours and more efficient fermentations (37, 38). Numerous recent studies have demonstrated how interspecific hybridization can be applied to increasing both lager yeast diversity and fermentation performance (1–3, 32). This ‘natural approach’ is a particularly attractive strain development tool for the brewing industry, because the use of genetically modified yeast is still not common as a result of regulations and public opinion (39). Another such GM-free strain development tool is adaptive evolution, which has also been successfully applied to improve several brewing-relevant traits in yeast (14, 17, 18, 20, 21, 29). Here we demonstrate how adaptive evolution can be applied to newly created interspecific lager yeast hybrids, in order to further improve their fermentation traits, and reveal the genetic changes that have occurred in the variant strains during adaptation.

By performing 30 consecutive batch fermentations in media supplemented with 10% ethanol, we aimed to generate and select ethanol-tolerant variants of four different brewing yeast strains; 3 of which were interspecific lager hybrids between *S. cerevisiae* and *S. eubayanus*. While experimental evolution is typically carried out in chemo- or turbidostats to allow for constant growth in defined nutrient availability (40), we here chose to use serial batch cultures for simplicity and to mimic the growth cycle the yeast encounters in repeated use in brewery fermentations (10). Our results show that the amount of yeast produced during each one-week fermentation cycle positively correlated with the number of consecutive batch fermentations, indicating that the strains adapted to the high ethanol concentration in the growth media. Here, approximately 130 to 160 yeast generations were achieved with 30 batch fermentations. Previous studies on adaptive evolution for ethanol tolerance have shown an increase can be achieved after 140 to 480 generations (14, 17–19), with evidence of increased fitness already after 40 generations in media containing ethanol (14). As the yeast is not constantly in exponential growth, it is expected that the batch fermentation process used here is more time-consuming than a continuous setup, where similar results have been achieved in less time (19).

A two-step screening process was used to ensure that variants exhibiting improved fermentation both in wort and in the presence of ethanol were selected from the adapted population. While growth in the presence of ethanol has been shown to be weakly positively correlated with ethanol production (41), we chose to monitor and select based on sugar consumption instead of growth, since we were interested in improving fermentation. As was revealed from the initial high-throughput screening, the majority of the strains that were isolated throughout the adaptation process outperformed the wild-type strains in regards to consumption of maltose and maltotriose in the presence of ethanol. However, variants arising from experimental evolution may in some cases exhibit antagonistic pleiotropy, where evolved variants show better fitness only in the environment in which they were selected (14). To prevent this, we performed a final screening step in small-scale wort fermentations. As was revealed during these small-scale fermentations, several isolates did in fact perform worse than the wild-type strains in wort, despite outperforming the wild-type strains in the ethanol-containing media used during high-throughput screenings.

The 2L-scale fermentations revealed that all eight of the tested variant strains outperformed the wild-type strains from which they were derived. The exact mechanisms for this improvement were not elucidated, but results seem to suggest that both improved ethanol tolerance and maltose use could have contributed, particularly as differences between variant and wild-type strains seemed to appear as fermentation progressed. Results revealed that many of the variants exhibited improved ethanol tolerance and accumulation capacity, while isolates showed considerable improvements in maltose consumption during high-throughput screening and wort fermentation. In previous studies, where brewing strains have been adapted to very high-gravity wort conditions, variant strains have exhibited increased expression of α-glucoside transporters and genes involved in amino acid synthesis (20, 21). Genome analysis of ethanol-tolerant variants has revealed that ethanol tolerance is a complex process, affected by several different mechanisms, including general stress response, intracellular signalling, and cell wall and membrane composition and organization (14, 19, 42). Here, several changes in the genomes of the variant strains were observed that could potentially contribute to the improved fermentation performance and ethanol tolerance. No SNPs, structural variations or gene-level copy number changes were observed for the genes encoding α-glucoside transporters in the variant strains. However, whole-chromosome copy number gains of the *S. cerevisiae*-derived chromosome VII, containing *MAL31* and *AGT1*, were observed in several variants. Interestingly, non-synonymous mutations in *IRA2* were observed in three of the variant strains. This gene negatively regulates the RAS-cAMP pathway (36), which in turn is involved in regulating metabolism, cell cycle and stress resistance (43, 44). Adaptive mutations in this gene have been reported previously (45) after experimental evolution in glucose-limited media, with *ira2* deletion strains exhibiting increased fitness. Mutations in *IRA2* have also been reported for strains evolved for increased xylose fermentation (46).

In regards to ethanol tolerance, we only identified non-synonymous mutations in one gene, *UTH1*, that has previously been reported to enhance ethanol tolerance. In turbidostat evolution experiments in high-ethanol media, Avrahami-Moyal et al. (19) found mutations in *UTH1* in a fraction of the evolved clones, and showed that deletion of this gene enhanced ethanol tolerance. While testing the direct effect of the non-synonymous mutations listed in Table 2 on ethanol tolerance by reverse engineering was outside the scope of this particular study, we feel it would be valuable to confirm their role in response to ethanol stress. In fact, several of the genes that were affected here (*BST1*, *CBP1*, *DAL81*, *EAP1*, *HAP4*, *HSP150*, *IRA2*, *MHP1*, *RAT1*, *RKM3*, *SFL1*, *TOD6* and *YIM1*), were also found to contain mutations in the evolved clones that were isolated by Voordeckers et al. (14) following exposure to increasing ethanol concentrations. In addition to SNPs and Indels, copy number variations are commonly reported in adapted strains (14, 15, 47). While it is thought that chromosome copy number changes allow for a rapid route of adaptation, they have an non-specific effect on the phenotype (14). Here we observed several common chromosome losses and gains. The *S. cerevisiae*-derived chromosome XIV was amplified in several of the variant strains, and interestingly, this chromosome has been reported through QTL mapping to carry genes (*MKT1*, *SWS2*, *APJ1*) associated with increased ethanol tolerance (42).

The variants did not only ferment faster, but in most cases also produced higher amounts of desirable esters and lower amounts of unwanted off-flavours compared to the wild-type strains. This was unexpected, as we only selected for fermentation and genetic hitchhiking is common during adaptive evolution (33). In previous studies on brewing yeasts adapted to high-gravity conditions, Ekberg et al. (21) reported increased concentrations of unwanted diacetyl, while Blieck et al. (20) observed slight increases in higher alcohol and diacetyl concentrations. As the aroma profile was not monitored during the screening process, it is vital to ensure that it is satisfactory for any selected variants. Genome analysis did not reveal any obvious causes for the increase in ester formation and decrease in diacetyl formation, as no SNPs, Indels or gene-level copy number changes affected genes that have previously been reported to be linked with the formation of these compounds. Some genes, such as *ATF2* on chromosome VII and *ILV6* on chromosome III, were affected by chromosome-level copy number changes, and could therefore have altered expression levels. Furthermore, most of the observed mutations were intergenic, and they could therefore have an indirect effect on these phenotypes by affecting gene regulation. Loss of heterozygosity has also been reported to be a method of adaptation in hybrid strains (28), and here, for example, we observed loss of heterozygosity on the right arm of the *S. cerevisiae*-derived chromosome XV in multiple variant strains. This particular region contains *ATF1*, the gene encoding the main alcohol acetyltransferase responsible for acetate ester synthesis (48, 49), which in the *S. cerevisiae* A81062 genome contains four heterozygous SNPs, one of which is non-synonymous. The two alleles of *ATF1* may therefore have slightly different functionality.

Interestingly, the greatest improvements in fermentation compared to the wild-type strains were observed with the polyploid interspecific hybrids. An increased ploidy level may allow for more rapid adaptation (12, 15, 30), presumably from gaining beneficial mutations at higher rates, along with chromosome losses and aneuploidy. The genome size of both of the variants derived from the tetraploid hybrid Y2 had decreased, while it had increased slightly for those derived from the triploid hybrid Y3. Aneuploidy and convergence towards a diploid state has commonly been reported during evolutionary engineering (11, 14, 15). Surprisingly, the largest change in genome size was observed for one of the variants derived from the diploid *S. cerevisiae* parent size. The results indicate that under these adaptation conditions, it was not only hybrid strains that possessed an unstable genome, and adapted variants from an industrial ale strain could be obtained without any prior mutagenesis. Evolutionary engineering studies involving interspecific hybrids, have indicated that in certain conditions either of the parental sub-genomes may be preferentially retained depending on the selective pressure (5, 8, 28), while the other may be lost. Piotrowski et al. (5), for example, showed that growing *S. cerevisiae* × *S. uvarum* hybrids in high temperatures, resulted in the loss of the ‘heat-sensitive’ *S. uvarum* sub-genome. Here, we saw a greater loss of the *S. eubayanus* sub-genome in the variants derived from interspecific hybrids. It is therefore tempting to speculate that repeating the adaptation process at a lower temperature would have retained more of the *S. eubayanus* sub-genome in the variants, and this could be a target for future studies. In fact, the natural lager yeast hybrids of Saaz-type have retained a larger fraction of the *S. eubayanus* sub-genome compared to the *S. cerevisiae* sub-genome (50, 51), and it is still unclear whether exposure to cold temperatures have had any effect on its evolution.

In conclusion, adaptive evolution in high-ethanol media was successfully used to generate stable and superior variant strains from 4 different brewing strains, 3 of which were de novo interspecific lager yeast hybrids. These adapted variants outperformed the strains which they were derived from during wort fermentation, and the majority also possessed several desirable brewing-relevant traits, such as increased ester formation and ethanol tolerance, and decreased diacetyl formation. While not tested here, it is likely that many of the adapted variant strains would also outperform the wild-type strains in very high-gravity wort, i.e. wort containing over 250 g extract L^−1^, as these fermentations require good tolerance towards both high osmotic pressure and ethanol concentrations (10), which several of the variant strains demonstrated by their ethanol accumulation capacity. Our study demonstrates the possibility of improving *de novo* lager yeast hybrids through adaptive evolution, and these superior and stable variants are viable candidates for industrial lager beer fermentation.

## Materials & Methods

### Yeast strains

A list of strains used in this study can be found in Table 1. Three different *de novo* lager yeast hybrids, generated in previous studies by our lab (3, 32), along with a *S. cerevisiae* ale parent strain (common to all three hybrids) were subjected to the adaptation process. Eight variant strains (two from each of the four wild-type strains) were isolated and subjected to phenotypic and genetic analysis. The ploidy of all the strains was determined by flow cytometry as described previously (32).

### Adaptation in a high-ethanol environment

The adaptation process was carried out in batch fermentations to mimic consecutive industrial brewery fermentations. Yeasts were grown in sterile 2 mL screw-cap microcentrifuge tubes (VWR Catalog Number 10025-754) containing 1 mL of growth media. Four different yeast strains (Y1, Y2, Y3 and Y4) were used for the adaptation experiment (see Table 1 for more information). These were grown in two different adaptation media: M1 (1% yeast extract, 2% peptone, 2% maltose, 10% ethanol) and M6 (1% yeast extract, 2% peptone, 1% maltose, 1% maltotriose, 10% ethanol). Each batch fermentation was inoculated to a starting OD600 of 0.1 with yeast from the previous batch fermentation. The first batch fermentations were inoculated from pre-cultures that were grown overnight in YPM media (1% yeast extract, 2% peptone, 2% maltose). Tubes were incubated statically for 7 days at 18 °C. Three replicate tubes or adaptation lines (A, B, and C) were used for each yeast strain and media (A and B were never mixed). In order to avoid contamination, the optical density at the end of each batch fermentation was measured only from the third replicate (C), which was subsequently discarded following the OD600 measurement. After 10, 20 and 30 consecutive batch fermentations, 10 μL aliquots of the cell populations were spread onto agar plates containing solidified versions of the growth media (2% agar added) for isolation of variants showing rapid growth. The agar plates were incubated at 18 °C until colonies started emerging, and the two largest colonies from each plate were selected for further screening (for a total of four isolates per yeast strain, per media, per isolation time point). An overview of the adaptation process and initial isolation step is depicted in Figure 1A and 1B, respectively.

### Screening

The isolates were initially screened on 96-well plates using a Beckman Coulter liquid handling robot to select for fast fermenting variants. Strains were grown in Nunc™ 96-well polystyrene round bottom microwell plates (Thermo Scientific 268200), in 150 μl volume at 14 °C, with 1200 rpm agitation in a Thermo Scientific Cytomat Plate Hotel (1 mm throw). Pre-cultures were prepared by inoculating 10 μL aliquots of cell suspension from frozen stocks into 140 μL of media consisting of 6.2% malt extract (Senson Oy, Finland) in the plates. Pre-cultures were incubated for 4 days until all strains had reached stationary phase. The pre-culture plates were centrifuged and pellets were resuspended in 50 mM citrate buffer (pH 7.2) to deflocculate the yeast. 10 μL aliquots of these suspensions were used to inoculate 140 μL of screening media for the experimental cultures. The isolates were grown in a screening media consisting of 6.2% malt extract (Senson Oy, Finland), 5% ethanol and 10% sorbitol. The extract content of this media was approximately 5 °P (50 g/L). The ethanol was added to the screening media to replicate the conditions the yeast is exposed to towards the end of brewery fermentations, while the sorbitol was added to replicate the increased osmotic pressure the yeast is exposed to in the beginning of brewery fermentations when sugar-rich wort is used. Each isolate was grown in triplicate, while wild-type strains were grown in at least 12 replicates. Strains and replicates were distributed randomly on the 96-well plates. The fermentations were monitored by measuring the optical density at 595 nm every 3 hours using the DTX 880 multimode detector (Beckman Coulter) associated with the robot, and by drawing samples for HPLC analysis after 48, 96 and 144 hours. This screening step is depicted in Figure 1C. Three isolates per yeast strain and media (for a total of 24 isolates) were selected for further screening in small-scale wort fermentations based on the following criteria: 1) the highest sugar consumption after 144 hours, 2) the isolates must be from separate adaptation lines and isolation time points.

To ensure that the isolates were also able to ferment actual wort efficiently, a final screening step was conducted by carrying out a set of small-scale wort fermentations. The small-scale fermentations were carried out in plastic 50 mL centrifuge tubes capped with a glycerol-filled airlock. The 24 isolates selected from the previous screening step and the 4 wild-type strains were grown overnight in 50 mL YPM at 18 °C. The pre-cultured yeast was then inoculated into 30 mL of 15 °P all-malt wort at a rate of 15 × 10^6^ viable cells mL^−1^. Fermentations were carried out in duplicate at 15 °C for 9 days, and these were monitored daily by mass lost as CO_2_. This screening step is depicted in Figure 1D. The maximum fermentation rate of each strain was determined and one isolate per yeast strain and media (for a total of 8 isolates) were selected based on the following criteria: 1) the highest fermentation rate, 2) isolated after a larger number of batch fermentations. These eight isolates are listed in Table 1, and were further characterized in 2L-scale wort fermentations.

### 2L-scale wort fermentations

The eight variant strains were characterized in fermentations performed in a 15 °Plato high gravity wort at 15 °C. Yeast was propagated essentially as described previously (3), with the use of a ‘Generation 0′ fermentation prior to the actual experimental fermentations. The experimental fermentations were carried out in duplicate, in 2-L cylindroconical stainless steel fermenting vessels, containing 1.5 L of wort medium. The 15 °Plato wort (69 g maltose, 17.4 g maltotriose, 15.1 g glucose, and 5.0 g fructose per litre) was produced at the VTT Pilot Brewery from barley malt. Yeast was inoculated at a rate of 15 × 10^6^ viable cells mL^−1^. The wort was oxygenated to 15 mg L^−1^ prior to pitching (Oxygen Indicator Model 26073 and Sensor 21158, Orbisphere Laboratories, Switzerland). The fermentations were carried out at 15 °C until an apparent attenuation of 80% (corresponding to approx 7% alcohol (v/v)) was reached, or for a maximum of 14 days. Wort samples were drawn regularly from the fermentation vessels aseptically, and placed directly on ice, after which the yeast was separated from the fermenting wort by centrifugation (9000 × g, 10 min, 1 °C). Samples for yeast-derived flavour compounds analysis were drawn from the beer when fermentations were ended.

### Chemical analysis

Concentrations of fermentable sugars (maltose and maltotriose) were measured by HPLC using a Waters 2695 Separation Module and Waters System Interphase Module liquid chromatograph coupled with a Waters 2414 differential refractometer (Waters Co., Milford, MA, USA). A Rezex RFQ-Fast Acid H+ (8%) LC Column (100 × 7.8 mm, Phenomenex, USA) was equilibrated with 5 mM H_2_SO_4_ (Titrisol, Merck, Germany) in water at 80 °C and samples were eluted with 5 mM H_2_SO_4_ in water at a 0.8 mL min^−1^ flow rate.

The alcohol level (% v/v) of samples was determined from the centrifuged and degassed fermentation samples using an Anton Paar Density Meter DMA 5000 M with Alcolyzer Beer ME and pH ME modules (Anton Paar GmbH, Austria).

Yeast-derived higher alcohols and esters were determined by headspace gas chromatography with flame ionization detector (HS-GC-FID) analysis. 4 mL samples were filtered (0.45 μm), incubated at 60 °C for 30 min and then 1 mL of gas phase was injected (split mode; 225 °C; split flow of 30 mL min^−1^) into a gas chromatograph equipped with an FID detector and headspace autosampler (Agilent 7890 Series; Palo Alto, CA, USA). Analytes were separated on a HP-5 capillary column (50 m × 320 μm × 1.05 μm column, Agilent, USA). The carrier gas was helium (constant flow of 1.4 mL min^−1^). The temperature program was 50 °C for 3 min, 10 °C min^−1^ to 100 °C, 5 °C min^∑1^ to 140 °C, 15 °C min^−1^ to 260 °C and then isothermal for 1 min. Compounds were identified by comparison with authentic standards and were quantified using standard curves. 1-Butanol was used as internal standard.

Total diacetyl (free and acetohydroxy acid form) was measured according to Analytica-EBC method 9.10 (52). Samples were heated to 60 °C and kept at this temperature for 90 min. Heating to 60 °C results in the conversion of α-acetolactate to diacetyl. The samples were then analyzed by headspace gas chromatography using a gas chromatograph equipped with a μECD detector and headspace autosampler (Agilent 7890 Series; Palo Alto, CA, USA). Analytes were separated on a HP-5 capillary column (50 m × 320 μm × 1.05 μm column; Agilent, USA). 2,3-Hexanedione was used as an internal standard.

### Ethanol tolerance and accumulation capacity

As several of the wild-type and variant strains flocculated strongly, we were unable to reliably determine ethanol tolerance in liquid cultures based on optical density measurements. Therefore, we assessed ethanol tolerance based on the ability to grow on YPD agar plates supplemented with various levels of ethanol. Overnight pre-cultures of all the strains were grown in YPM at 25 °C. The yeast was then pelleted and resuspended in 50 mM citrate buffer (pH 7.2) to deflocculate the yeast. The cell concentration was measured with a Nucleocounter^®^ YC-100™ (ChemoMetec, Denmark), after which suspensions were diluted to contain approximately 10^5^, 10^4^ and 10^3^ cells mL^−1^. 5 μL aliquots of the suspensions of each strain was spotted onto agar plates containing YPD supplemented with 9%, 11% and 13% ethanol. Plates were sealed with parafilm, placed in ziplock bags, and incubated at 25 °C for up to 21 days.

The ethanol accumulation capacity of the strains was also assessed as described by Gallone et al. (53) with modifications. Overnight pre-cultures of all the strains were grown in YP-4% Maltose at 25 °C. The yeast was then pelleted and resuspended to an OD600 of 20 in 50 mM citrate buffer (pH 7.2) to deflocculate the yeast. 35 mL of YP-35% Maltose was then inoculated with the yeast strains to an initial OD600 of 0.5. Fermentations took place in 100 mL Erlenmeyer flasks capped with glycerol-filled airlocks. Flasks were incubated at 18 °C with gentle shaking (100 rpm) for 28 days. The mass loss was monitored to estimate when fermentation finished. After the fermentations had finished, the cultures were centrifuged, after which the alcohol content of the supernatants was measured with an Anton Paar Density Meter DMA 5000 M with Alcolyzer Beer ME and pH ME modules (Anton Paar GmbH, Austria).

### Genetic stability of variant strains

The genetic stability of the eight variant strains (Table 1) was assessed by culturing them repeatedly in YP-4% Maltose at 18 °C for over 80 generations (2, 9). After this, DNA was extracted from two randomly chosen isolates from each variant strain. DNA fingerprints were produced for each isolate and the eight variant strains with PCR using delta12 (5′-TCAACAATGGAATCCCAAC-3′) and delta21 (5′-CATCTTAACACCGTATATGA-3′) primers for interdelta DNA analysis (54). The DNA fingerprints of the isolates obtained after 80 generations were compared with those of the variant strains, and the variants were deemed genetically stable if the fingerprints were identical.

### Genome sequencing and analysis

Wild-type strains Y1 and Y2 have been sequenced in previous studies (3, 32), and reads for these strains were obtained from NCBI-SRA (SRX1423875 and SRX2459842, respectively). For this study, wild-type strains Y2-Y4 and the eight variant strains were sequenced by Biomedicum Genomics (Helsinki, Finland). In brief, DNA was initially isolated using Qiagen 100/G Genomic tips (Qiagen, Netherlands), after which an Illumina TruSeq LT pair-end 150 bp library was prepared for each strain and sequencing was carried out with a NextSeq500 instrument. Pair-end reads from the NextSeq500 sequencing were quality-analysed with FastQC (55) and trimmed and filtered with Cutadapt (56). Alignment of reads was carried out using SpeedSeq (57). Reads of *S. cerevisiae* Y1 (VTT-A81062) and its variants were aligned to a previously assembled reference genome (available under BioProject PRJNA301545) of the strain (3), while reads of hybrid strains Y2-Y4 and their variants were aligned to concatenated reference sequences of *S. cerevisiae* VTT-A81062 and *S. eubayanus* FM1318 (58) as described previously (3). Quality of alignments was assessed with QualiMap (59). Variant analysis was performed on aligned reads using FreeBayes (60). Variants in wild-type and variant strains were called simultaneously (multi-sample). Prior to variant analysis, alignments were filtered to a minimum MAPQ of 50 with SAMtools (61). Structural variation analysis was performed with LUMPY (62), Manta (63), and Scalpel (64). Variants that were unique to the variant strains (i.e. not present in the wild-type strain) were obtained with SnpSift (65). Annotation and effect prediction of the variants was performed with SnpEff (66). The filtered and annotated variants were finally manually inspected in IGV (67). Copy number variations were estimated based on coverage with CNVKit (68). The median coverage over 10,000 bp windows was calculated with BEDTools (69).

### Data visualization and analysis

Data and statistical analyses were performed with R (http://www.r-project.org/). Flow cytometry data was analysed with ‘flowCore’ (70) and ‘mixtools’ (71) packages. Growth curves from the high-throughput screening cultivations were produced based on optical density measurements using the logistic model in the ‘grofit’ package (72). Scatter, box and heatmap plots were produced in R. The ‘Circos-like’ plots in Figures S4-S15 in the Supplementary material were produced with the ‘circlize’ package (73). Significance between variant wild-type strains was tested by Student’s t-test (two-tailed, unpaired, and unequal variances).

### Data availability

The Illumina reads generated in this study have been submitted to NCBI-SRA under BioProject number PRJNA408119 in the NCBI BioProject database (https://www.ncbi.nlm.nih.gov/bioproject/).

## Acknowledgements

We thank Sue James for valuable comments during the study, Virve Vidgren for performing DNA extractions, Eero Mattila for wort preparation and other assistance in the VTT Pilot Brewery, and Aila Siltala for skilled technical assistance. This work was supported by the Alfred Kordelin Foundation, Svenska Kulturfonden - The Swedish Cultural Foundation in Finland, Suomen Kulttuurirahasto, SABMiller (ABInBev), and the Academy of Finland (Academy Project 276480).

